# Spacing theta-burst stimulation enhances synaptic potentiation in the vulnerable prefrontal cortex

**DOI:** 10.64898/2026.08.27.746330

**Authors:** Angela Zolis, Angel Hsin-Yun Hsieh, Sridevi Venkatesan, Rachael Ingram, John Georgiou, Christoph Zrenner, Graham L. Collingridge, Tarek K. Rajji, Evelyn K. Lambe

**Affiliations:** Department of Physiology, Temerty Faculty of Medicine, University of Toronto, Toronto ON, Canada; Hospital for Sick Children, Toronto ON, Canada; Lunenfeld-Tanenbaum Research Institute, Mount Sinai Hospital, Sinai Health System, Toronto ON, Canada; Tanz Centre for Research in Neurodegenerative Diseases, University of Toronto, Krembil Discovery Tower, Toronto, Ontario, Canada; Centre for Addiction and Mental Health (CAMH), Toronto ON, Canada; Department of Psychiatry, Temerty Faculty of Medicine, University of Toronto, Toronto ON, Canada; Department of Psychiatry, University of Texas Southwestern Medical Center, Dallas, Texas, USA; Department of OBGYN, Temerty Faculty of Medicine, University of Toronto, Toronto ON, Canada

## Abstract

Neuromodulation with intermittent theta-burst stimulation (iTBS) is a clinical treatment for major depression. One postulated mechanism of iTBS is to strengthen the synaptic connections which activate the prefrontal cortex to regulate mood. With *ex vivo* electrical stimulation and neuronal calcium imaging, we demonstrate that a common, clinical iTBS pattern (600-stimuli, ∼3-minutes) reliably strengthens the synaptic recruitment of the adult mouse prefrontal cortex. This synaptic potentiation, however, becomes less reliable in the depressive-like mouse model of prolonged social isolation. To better understand this change, we examine the impact of social isolation on the complex pattern of calcium elevation *during* clinical iTBS. In social isolation, the calcium peaks rise more prominently, but the induction peak no longer predicts the synaptic potentiation outcome. To better regulate calcium dynamics during induction, we test a paradigm with fewer iTBS episodes, separated by longer intervals. This spaced iTBS (90-stimuli, ∼10-minutes) in the adult prefrontal cortex limits calcium elevation during induction and yields reliable long-term potentiation (LTP) following either juvenile- or adult-onset social isolation. This research illustrates new strategies to interrogate and to enhance synaptic plasticity in the vulnerable prefrontal cortex.

**Significance Statement:** Neuromodulation therapy in major depression aims to boost synaptic potentiation in the prefrontal cortex. Since this region lacks simple behavioral readouts, preclinical studies using rodent brain slices are valuable to test and refine stimulus patterns. Here, we demonstrate that *ex vivo* neuronal calcium imaging reliably captures the strength and spatial spread of synaptic plasticity in the mouse prefrontal cortex in response to a common, clinical protocol (intermittent theta-burst stimulation, iTBS). This approach is sufficiently sensitive to detect problems in synaptic potentiation associated with social isolation and to identify translationally relevant changes to improve prefrontal plasticity in the vulnerable brain.

## Introduction

Major depression is increasingly treated with neuromodulation therapies applied to the prefrontal cortex (Chistyakov et al., 2010; Duprat et al., 2016; Li et al., 2014; Watson et al., 2023; Dalhuisen et al., 2024; Santos et al., 2026). A critical goal of these treatments is to strengthen the excitatory synapses (Di Lazzaro et al., 2008) that activate prefrontal neurons responsible for higher-order cognitive functions such as decision making, emotional processing and regulation, attention, and motivation (Friedman & Robbins, 2022; Szczepanski & Knight, 2014; Xu et al., 2019; Riddle et al., 2026). This treatment is necessary because there are profound disruptions in these functions in mood disorders, together with structural and neurophysiological abnormalities in the prefrontal cortex (Holmes et al., 2019; Keller et al., 2019; Liu et al., 2017; Pizzagalli & Roberts, 2022).

To date, the most effective and efficient neuromodulation pattern is intermittent theta-burst stimulation (iTBS) (Blumberger et al., 2018; Bulteau et al., 2022). This pattern administered through transcranial magnetic stimulation is effective in both treatment-resistant and first-episode depression (Voigt et al., 2019) and, overall, has a ∼40% patient response rate after multiple sessions of treatment (Blumberger et al., 2018; Bulteau et al., 2022; Dalhuisen et al., 2024; Santos et al., 2026). Such paradigms are administered to the prefrontal cortex and even one session (600 stimuli, 3-min) enhances synaptic recruitment in this brain region (Chung et al., 2017). For clinical treatment, the paradigm is typically repeated for multiple sessions/week for more than four weeks, for a total of ∼12,000 stimuli (Blumberger et al., 2018; Bulteau et al., 2022). Given the short duration of each treatment session (i.e. 3-minutes), clinical researchers have tested intensified versions of iTBS, prolonging the duration of each session, and repeating these sessions multiple times a day to give as many as 90,000 stimuli over several days (Cole et al., 2022; Cole et al., 2022; Ramos et al., 2025). While these accelerated iTBS protocols may bring a treatment response sooner (Hanlon et al., 2026; Tendler et al., 2026), there is ongoing debate about whether longer-lasting effects may be achieved by standard iTBS treatment. “More” brain stimulation is not necessarily better. Targeted refinements to the essential elements of iTBS may still be needed to improve overall treatment efficacy.

This growing interest in iTBS protocols has led to the first preclinical investigations of the synaptic impact of these protocols in mice *in vivo* (Gongwer et al., 2026; Johnson et al., 2026; Salimi et al., 2026). Experimental studies in the laboratory allows for controlled investigation of diverse models relevant to depression, including stress paradigms (Gongwer et al., 2026) and corticosterone treatments (Johnson et al., 2026). These studies apply the iTBS pattern using different modes of stimulus delivery, including transcranial magnetic stimulation, optogenetic stimulation, and electrical stimulation (Gongwer et al., 2026; Johnson et al., 2026; Salimi et al., 2026). They also permit a greater diversity of tools to measure prefrontal cortical plasticity.

*Ex vivo* experiments in brain slices offer additional opportunities to test local circuit impact and to refine iTBS protocols for greater impact. For context, theta-burst stimulation was first applied in human patients because *ex vivo* synaptic research demonstrated that this pattern of electrical stimulation elicits long-term potentiation (LTP) (Larson & Lynch, 1986; Bliss & Collingridge, 1993). Further refinements to theta-burst stimulation continue to be actively pursued *ex vivo* in hippocampal brain slices with promising results. Spaced theta-burst paradigms are of increasing interest. These paradigms change the architecture of the iTBS session, increasing the duration of the time between iTBS episodes. This extra time allows for the activation of second messengers to enhance the longer-lasting, protein-synthesis dependent LTP (Woo et al., 2003; Kim et al., 2010; Park et al., 2014, 2016, 2021).

Yet relatively little *ex vivo* research has investigated the current clinical iTBS pattern, nor pursued such stimulation in brain slices of the adult prefrontal cortex, let alone investigated newer concepts such as spaced iTBS. The results of experiments in the hippocampus, the brain region where most plasticity experiments are conducted, do not necessarily extrapolate to the prefrontal cortex, which is structurally and neurobiologically distinct (Carlén, 2017; Vogt et al., 2013; Vogt & Paxinos, 2014; Wang et al., 2008). In fact, many of the studies of theta-burst stimulation in the prefrontal cortex have had to suppress inhibition in order to achieve potentiation (Caballero et al., 2014; Hurtado et al., 2021; Martin et al., 2016; Thomazeau et al., 2014). Others have restricted their research to juvenile rodents (Kerkhofs et al., 2018; Lee et al., 2021, 2023), whose cortical inhibitory circuits are still maturing (Ferrer et al., 2018; Miyamae et al., 2017; Sydnor et al., 2026). Furthermore, it is important to note that clinical iTBS differs in episode and burst characteristics compared to protocols typically used in *ex vivo* research. To date, relatively little is known about the *ex vivo* synaptic impact of the clinical iTBS (600 stimuli, 3-min) in the adult prefrontal cortex, its mechanisms, and their vulnerability in rodent models with brain changes relevant to depression.

Here, we use *ex vivo* neuronal calcium imaging in response to electrical stimulation to demonstrate that clinical iTBS reliably strengthens prefrontal synapses in brain slices from adult group-housed mice. This reliability is decreased in brain slices from adult socially-isolated animals. The latter show prominent calcium elevations during iTBS, but these signals do not possess their typical power to predict the plasticity outcome. As a result, we aimed to refine the iTBS protocol to constrain calcium during induction. We tested a spaced iTBS protocol with fewer episodes and longer intervals between respective trains of high-frequency stimulation. Spaced iTBS (90 stimuli, 10-minutes) achieved stronger synaptic potentiation in socially-isolated mice than in group-housed controls, demonstrating that iTBS-elicited prefrontal plasticity is strongly influenced by the brain changes evoked by social isolation.

## Results

### Widefield neuronal calcium imaging to probe synaptic plasticity

To investigate iTBS-evoked synaptic plasticity in prefrontal cortex using widefield calcium imaging, we prepared brain slices from adult (> 70 days postnatal; **Table 1**) *Thy1*-GCaMP6f hemizygous mice of both sexes in equal proportions. Positioning the stimulating electrode in superficial cortex (**Figure 1A-B**), we established a baseline by imaging the response to test stimuli (5 stimuli of 100 µs duration given at 0.1 Hz; **Figure 1C-D**). The test stimulus was selected to match the pulse duration commonly used in transcranial magnetic stimulation (Spampinato et al., 2023). To test synaptic potentiation, brain slices received either clinical iTBS or a test pulse control (**Figure 1E-F**). We then followed the test pulse signals for 60 minutes to assess changes in their amplitude and spatial spread with respect to baseline.

**Figure 1:**
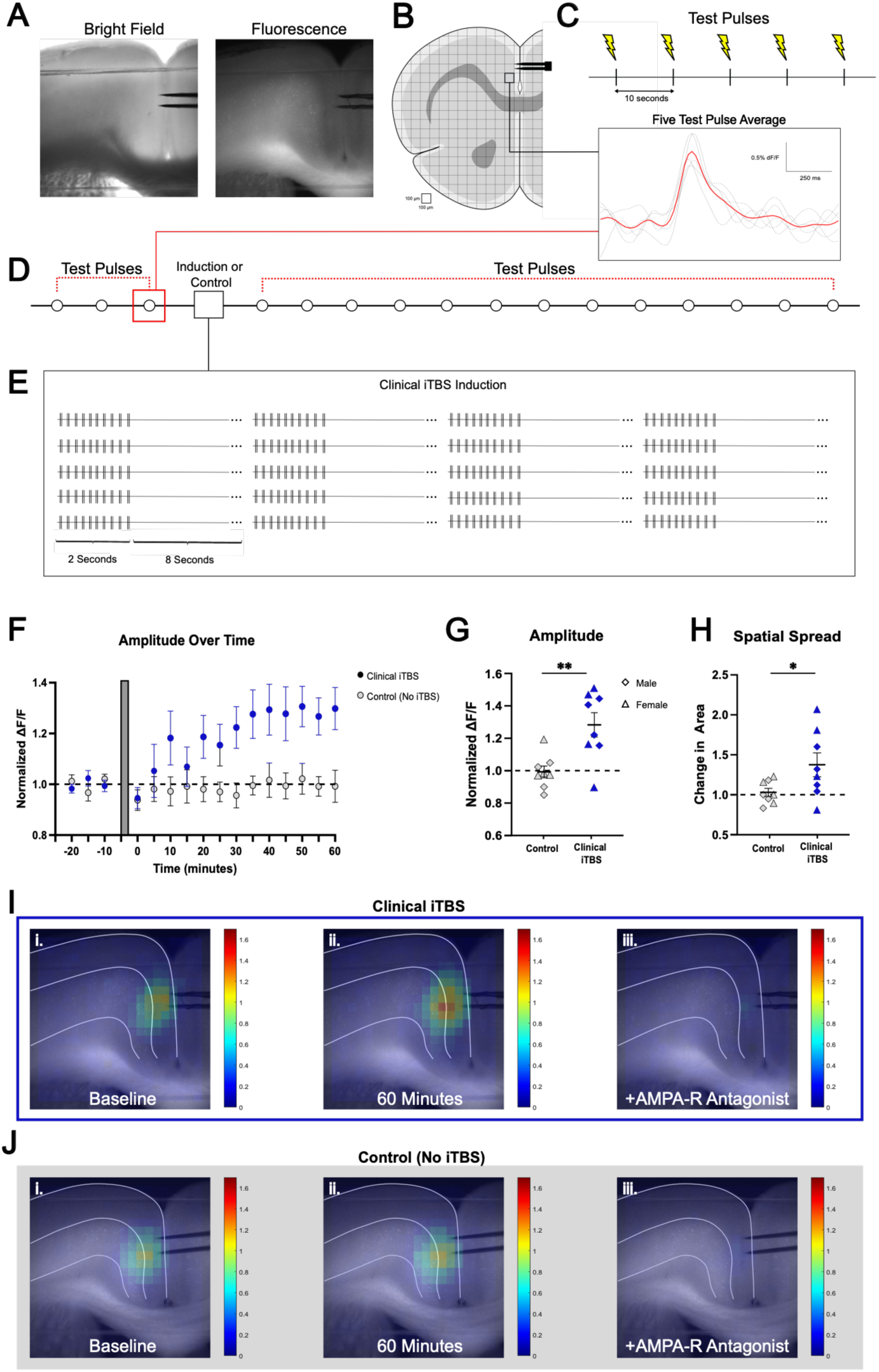
Impact of clinical intermittent theta-burst stimulation (iTBS) on long-term potentiation of neuronal calcium signals evoked by single test pulses. A. Prefrontal cortex brain slice under brightfield and fluorescence. B. Schematic of regions receiving test pulse stimulation while imaged for GCaMP6f calcium fluorescence. C. Schematic of test pulse protocol with individual (gray) and averaged (red) calcium signals. D. Schematic of the experimental approach with timing of imaged test pulse stimuli and clinical iTBS. E. Schematic of clinical iTBS protocol showing the 20 episodes of theta-burst stimulation. F. Time-course showing calcium changes (ΔF/F) upon test pulse delivery for control (gray circles) and clinical iTBS (blue circles). Gray bar denotes time of induction or control. G. Scattergram of calcium signal changes (Δ area) following control (no iTBS) or clinical iTBS induction. H. Scattergram of calcium signal spatial spread outcomes (for control versus clinical iTBS). Heatmaps (calcium level pseudocolour scale as shown) from (I) clinical iTBS or (J) control at (i) baseline, (ii) 60 minutes after induction or control, and (iii) after blockade of AMPA receptors. Data points shown as mean ± SEM; \**P* < 0.05, \*\**P* < 0.01.

**Table 1.** Age (mean ± SD, days postnatal) and housing condition of mice in the experiments for each Figure. Each group has equal numbers of male and female mice. For Figure 1, all mice were group housed. For Figures 2 and 4, mice were either group housed or socially isolated at weaning (∼P21) and isolation continued until brain slice imaging. Figure 3A-E includes the same mice as Figure 2, and Figure 3F-J includes the same mice as Figure 4. For juvenile-onset social isolation, mice were single housed at weaning until brain slice imaging (mean ± SD, 18 ± 5 weeks, range 10-23 weeks). For adult-onset social isolation, mice were single housed during adulthood (>60 days postnatal) until brain slice imaging (4 ± 4 weeks, range 1-15 weeks). The summary graphs at the bottom of Figures 4 and 5 are based on data from multiple figures.

| **Age Data** | **Control**<br>**Mean ± SD** | **Clinical iTBS**<br>**Mean ± SD** |
| --- | --- | --- |
| **Figure 1** | 122 ± 36<br>(8 mice) | 146 ± 34<br>(8 mice) |
| **Data** | **Group Housed**<br>**Mean ± SD** | **Socially Isolated**<br>**Mean ± SD** |
| **Figure 2** | 151 ± 37<br>(10 mice) | 150 ± 32<br>(12 mice) |
| **Figure 3A-E** | 151 ± 37<br>(10 mice) | 150 ± 32<br>(12 mice) |
| **Figure 3F-J** | 146 ± 33<br>(10 mice) | 148 ± 35<br>(10 mice) |
| **Figure 4** | 146 ± 33<br>(10 mice) | 148 ± 35<br>(10 mice) |
| **Figure 5** | n/a | 160 ± 54<br>(10 mice) |

Control stimulation experiments (no iTBS) showed that test pulses evoke consistent calcium signals in the prefrontal cortex across time (**Figure 1F-H**, one sample tests, amplitude: *t*(7) = 0.1992, *p* = 0.8478; spatial spread: *t*(7) = 0.6049, *p* = 0.5643). By contrast, clinical iTBS (20 episodes of 10 theta bursts, delivered every 10 s) potentiated the calcium signal at ∼60 minutes post-iTBS compared to baseline (one sample tests, amplitude: *t*(7) = 3.793, *p* = 0.0068; spatial spread: *t*(7) = 2.526, *p* = 0.0395). The outcome of clinical iTBS was significantly greater than control in terms of calcium signal amplitude (**Figure 1F-G**; unpaired *t*-test: *t*(14) = 3.499, *p* = 0.0035) and spatial spread (**Figure 1H**; unpaired *t*-test: *t*(14) = 2.195, *p* = 0.0455). We did not detect sex differences in the iTBS-induced potentiation (unpaired *t*-tests, amplitude: *t*(6) = 0.2977, *p* = 0.7760; spatial spread: *t*(6) = 0.6700, *p* = 0.5278). In both the clinical iTBS and control experiments, the synaptic nature of the test pulse signals was confirmed by their strong and significant suppression by the AMPA receptor antagonist NBQX (**Figure 1I-J**, right; paired *t*-tests amplitude: *t*(15) = 16.60, *p* < 0.0001; spatial spread: *t*(15) = 13.30, *p* <0.0001). Together, these experiments demonstrate that clinical iTBS significantly potentiates glutamatergic synapses of the major output neurons in the prefrontal cortex.

### Impact of social isolation on synaptic plasticity from clinical iTBS

We next investigated the impact of clinical iTBS on prefrontal synaptic plasticity in a socially-isolated vulnerable mouse model (Csikós et al., 2024; Grigoryan et al., 2022; Ieraci et al., 2016; Medendorp et al., 2018). With group-housed age- and sex-matched littermates as controls (**Figure 2A**), we first investigated evoked calcium responses and then the impact of clinical iTBS in adult mice of both sexes that had been socially isolated from weaning (**Table 1**). Investigating across a range of different strengths of test pulse stimulations, we did not detect differences in calcium responses between group housed and socially isolated mice (**Suppl Fig S1A**), and the selected test pulse currents did not differ with housing condition (**Suppl Fig S1B**; unpaired *t*-test; *t*(20) = 0.8016, *p* = 0.4322).

**Figure 2:**
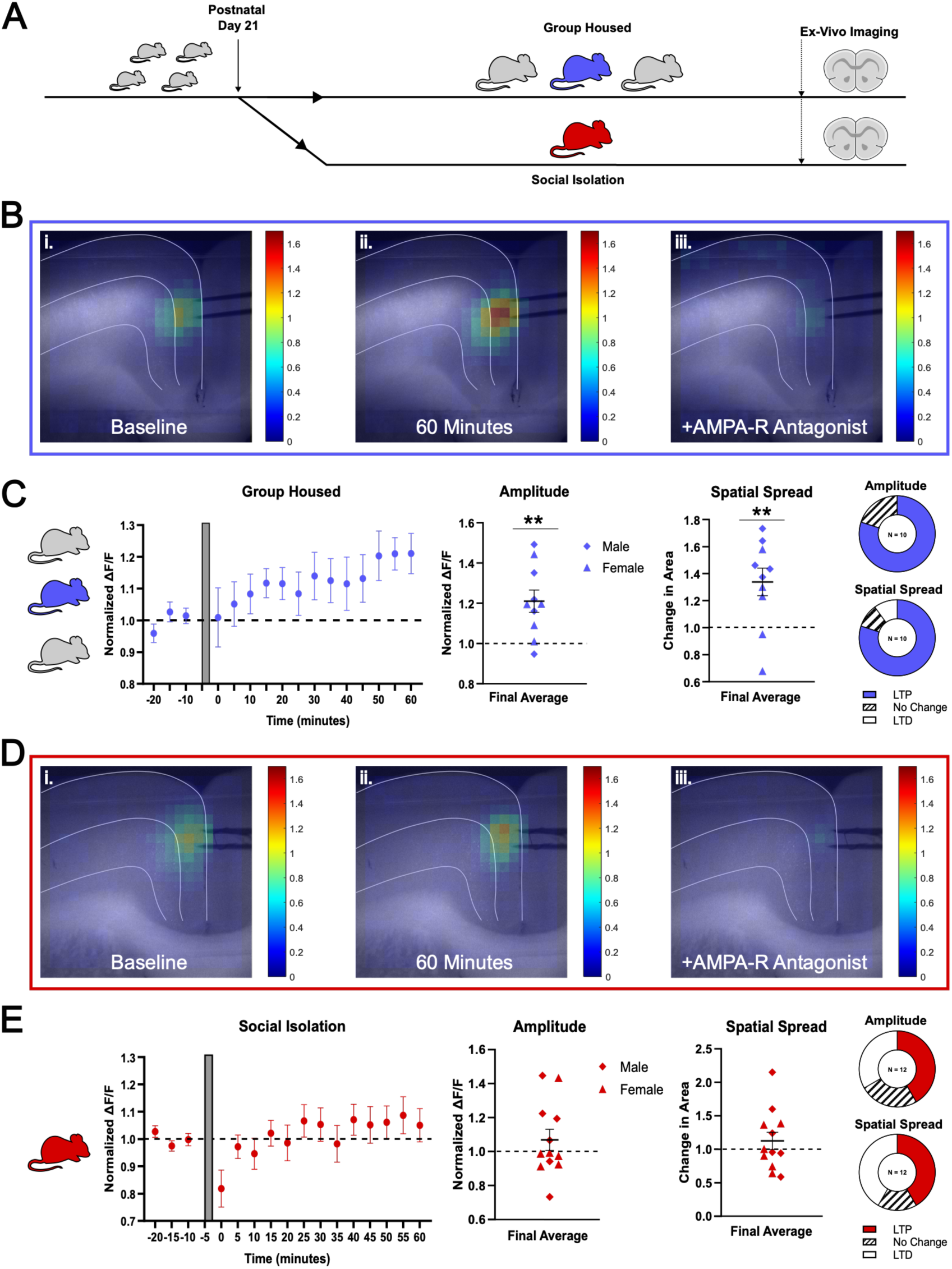
Social isolation influences clinical iTBS plasticity outcome. **A**. Schematic of social isolation from weaning (red mouse) and group-housed control (blue mouse) with timing of *ex vivo* experiments. **B**. Example calcium signal heatmaps from group-housed mice evoked by test pulses at (i) baseline, (ii) one hour after clinical iTBS, and (iii) after blockade of AMPA receptors. **C**. Time course showing test pulse amplitudes from group-housed mice (gray bar is clinical iTBS). Scattergrams of amplitude and spatial spread outcomes. Donut charts show proportions of different types of response (no change defined as ± 5%). **D**. Social isolation example heatmaps. **E**. Time course of amplitude outcomes from socially isolated mice; gray bar depicts clinical iTBS. Scattergrams of calcium signal amplitude and spatial spread outcomes, with donut charts showing proportions of different types of response. Mean ± SEM; \*\**P* < 0.01.

Again, clinical iTBS significantly potentiated the signals evoked by test pulses in the group-housed controls (**Figure 2B-C**; one sample tests; amplitude: *t*(9) = 3.762, *p =* 0.0045; spatial spread: *t*(9) = 3.324, *p =* 0.0089), with calcium responses significantly suppressed by AMPA receptor antagonist (amplitude: paired t-tests, *t*(9) = 11.46, *p <* 0.0001; spatial spread: paired t-test, *t*(9) = 16.00, *p <* 0.0001). In social isolation, however, clinical iTBS was less robust (**Figure 2D-E**). Neither the amplitude nor the spatial spread rose to the level of a significant increase in the socially isolated mice (**Figure 2E**; amplitude: one-sample t-tests, *t*(11) = 1.102, *p* = 0.2939; spatial spread: one-sample t-test*, t*(11) = 0.9723, *p* = 0.3518). In both groups, test pulse responses were significantly suppressed by NBQX (paired t-tests; amplitude: *t*(11) = 11.44, *p <* 0.0001; spatial spread: (*t*(11) = 5.561, *p* = 0.0002).

Combining the group-housed mice from Figures 1 and 2, clinical iTBS yielded significantly greater potentiation outcome as measured by calcium signal amplitude for group-housed mice compared to socially-isolated mice (paired *t*-test; *t*(28) = 2.324, *p* = 0.0276), with a trend observed for increased spatial spread (paired *t*-test; *t*(28) = 1.882, *p* = 0.0702). The prefrontal cortex from group-housed mice showed LTP in most brain slices; whereas the socially-isolated mice showed outcomes that included a substantial proportion with LTD.

### Impact of social isolation on calcium responses during theta-burst stimulation

To probe calcium dynamics *during* clinical iTBS, we examined the calcium responses across the 20 episodes (also called trains) of theta-burst induction. The first episode shows a relatively constrained calcium response. Subsequent episodes rise further, plateau, and subsequently decline (**Figure 3A**). Area-under-the-curve (AUC) measures reveal a significant interaction between induction episode responses and housing conditions (**Figure 3B**; repeated two-way ANOVA with Geisser-Greenhouse correction, *F*(2.044, 40.89) = 3.607; *p* = 0.0352), with significantly greater responses detected in episode 2-7 (Dunnett *post hoc* tests, *p* < 0.05). Socially-isolated mice show a significantly steeper rising slope from first to peak episode compared to group-housed mice (**Figure 3C-D**; unpaired t-test, *t*(20) = 2.202, *p* = 0.0400).

**Figure 3:**
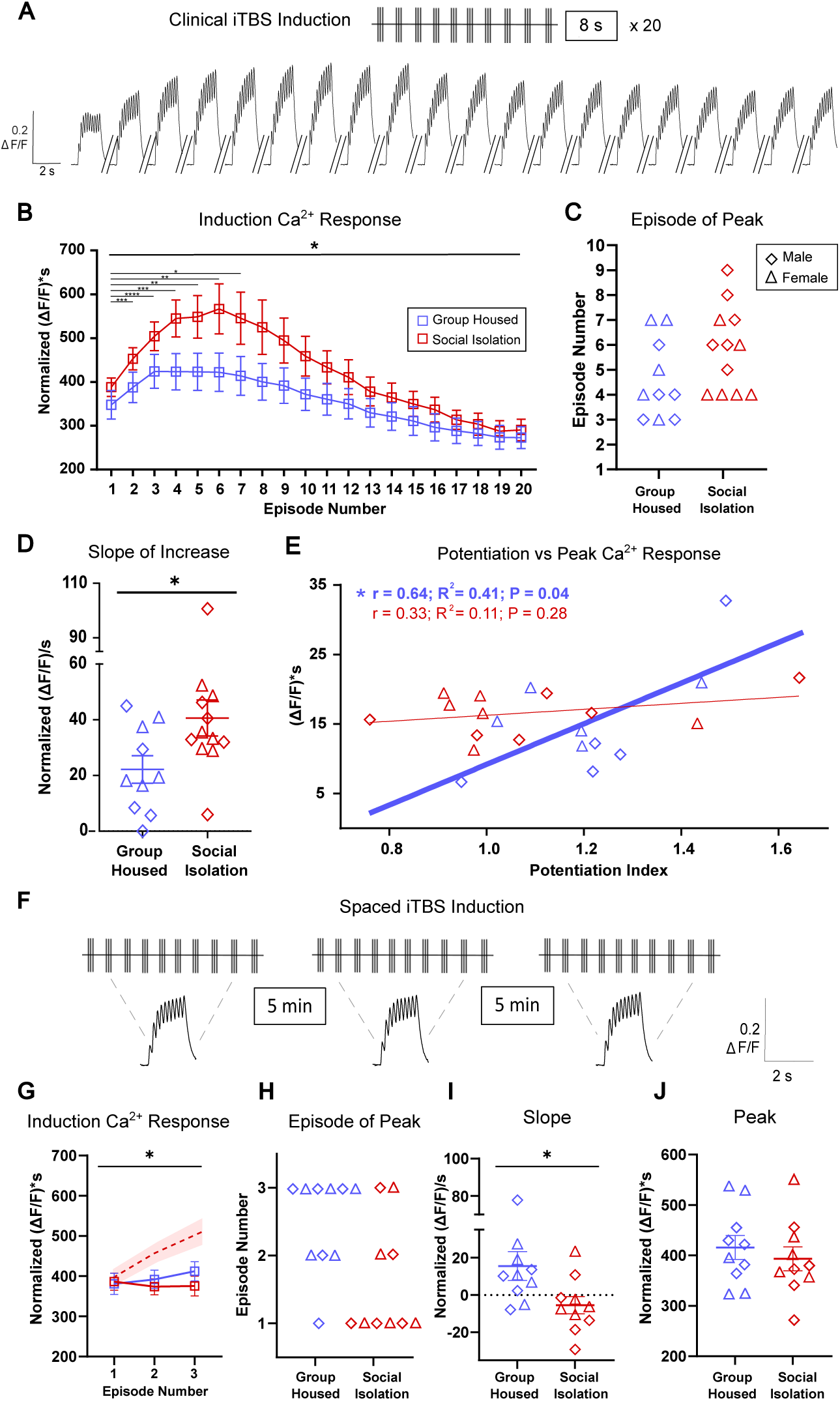
Aberrant elevation of calcium during clinical iTBS in social isolation is prevented by *spaced* iTBS. **A**. Schematic of clinical iTBS and example of per-episode calcium responses. **B**. Graph of the per-episode calcium elevations (area under curve ([ΔF/F]*s) normalized to test pulse response) during clinical iTBS induction in group-housed (blue) and socially-isolated mice (red). **C**. Scattergram showing which episodes had the peak calcium elevations across the mice. for clinical iTBS. **D**. Scattergram showing calcium signal slopes from first to peak episode ([ΔF/F]/s). **E**. Correlation plots of peak calcium response ([ΔF/F]*s) versus test pulse-evoked potentiation outcome for group-housed and socially-isolated mice. **F**. Schematic of spaced iTBS and example of per-episode calcium responses. **G**. Graph of per-episode calcium elevations during spaced iTBS induction (dashed line contrasts the trajectory of the first 3 episodes from clinical iTBS). **H**. Scattergram showing which episodes had peak calcium responses in spaced iTBS. **I**. Scattergram of slopes across spaced iTBS. **J**. Scattergram of peak calcium response during spaced iTBS. Mean ± SEM; \**P* < 0.05,** *P <* 0.01, \*\*\**P*<0.001, \*\*\*\**P <* 0.0001.

Since calcium increases are considered essential to strengthen neuronal connections (Shouval et al., 2002; Yeung et al., 2004; Nishiyama et al., 2000), we examined the relationship between the peak calcium response *during* clinical iTBS, and the final synaptic plasticity achieved. As hypothesized, the peak-episode calcium measure correlated significantly with the potentiation outcome in group-housed mice (**Figure 3E**; R^2^ = 0.42, *p* = 0.044), a relationship that holds if the group-housed mice from Figures 1 and 2 are combined (R^2^ = 0.27, *p* = 0.027). By contrast, this association was not detected in social isolation (R^2^ = 0.11, *p* = 0.284). Intriguingly, the iTBS calcium measures are strong in socially isolated mice, but their peaks appear dissociated from the plasticity outcome. This discrepancy raises the question whether calcium during theta-burst stimulation can be detrimental for potentiation under some circumstances.

### Theta-burst induction can be altered to constrain calcium responses

Since the calcium measures from the first episode of iTBS are similar across group-housed and isolated conditions (unpaired *t*-test *t*(20) = 1.077; *p* = 0.2944), we hypothesized that a spaced iTBS protocol, with fewer iTBS episodes (3 vs 20 episodes) separated by longer intervals (every 5 minutes vs every 10 s), would constrain calcium elevation during iTBS (**Figure 3A and F)**. Such spaced induction approaches have been successfully used to generate long lasting synaptic plasticity in *ex vivo* hippocampal experiments (Woo et al., 2003; Kim et al., 2010; Park et al., 2014, 2016, 2018, 2021; Pauli et al., 2025). However, this prior hippocampal research employed different burst characteristics than clinical iTBS (i.e. 5 stimuli at 100 Hz instead of the 3 stimuli at 50 Hz). Here, we deliver 3 episodes that are identical to the episodes in clinical iTBS, except for their 5-minute inter-episode intervals. That is, instead of applying the 600-stimuli protocol used in clinical iTBS (**Figure 3A**), we deliver only 90 pulses (**Figure 3F**).

We initially evaluated spaced iTBS to compare calcium responses in the group-housed versus isolated mice (**Figure 3F-G**, **Table 1**). As hypothesized, the calcium measures elicited by spaced iTBS were constrained. There was a significant interaction between episode measure and housing condition (**Figure 3G**; repeated two-way ANOVA with Geisser-Greenhouse correction, *F*(1.573, 28.32) = 4.024, *p* = 0.0376). There were small differences in spaced iTBS induction responses between group-housed and socially-isolated mice, specifically their peaks occur in different episodes (**Figure 3H**) and their slopes differ significantly (**Figure 3I**; unpaired t-test, *t*(18) = 2.336, *p* = 0.0312). However, we did not detect a difference in peak calcium response during spaced iTBS between group-housed and socially-isolated mice (**Figure 3J**; unpaired t-test, *t*(18) = 0.67, *p* = 0.51).

### Impact of spaced iTBS on synaptic plasticity and comparison to clinical iTBS

Since spaced iTBS was effective at controlling calcium levels during induction in socially-isolated mice, we probed its impact on synaptic plasticity. To investigate the impact of spaced iTBS on prefrontal synaptic plasticity, we recorded the calcium signal evoked by the test pulse following induction. Socially-isolated mice showed significant increases in both amplitude (**Figure 4A-B**, one-sample test, *t*(9) = 2.850, *p* = 0.0191) and spatial spread of the test pulse signal (*t*(9) = 3.344, *p =* 0.0086), suggesting that spaced iTBS is effective at inducing synaptic plasticity in socially-isolated mice.

**Figure 4:**
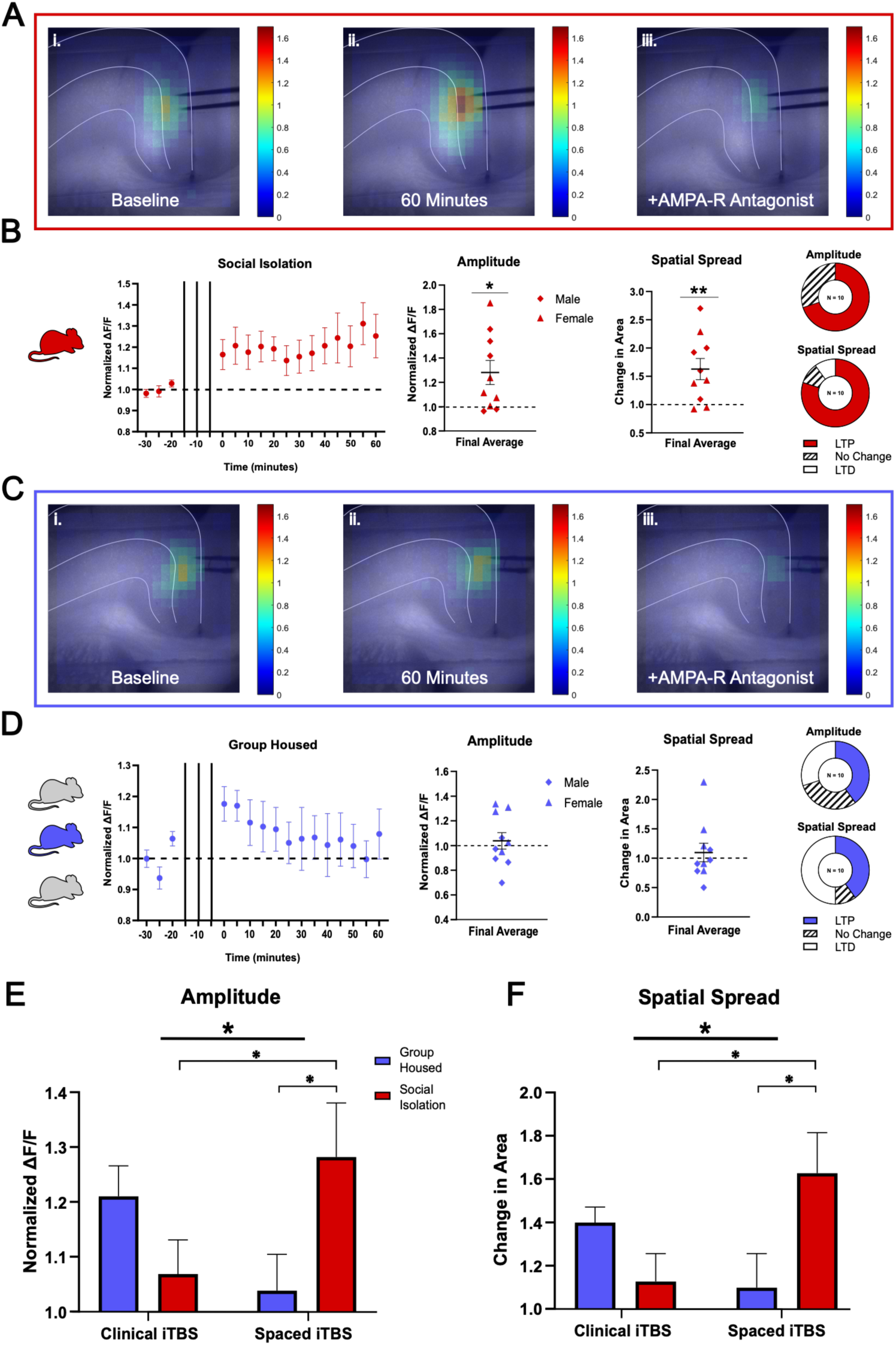
Potentiation via spaced iTBS is effective in socially-isolated mice. **(A)** Example heatmaps from socially isolation with test pulse signal at (i) baseline, (ii) one hour after clinical iTBS, and (iii) after blockade of AMPA receptors. **B.** Time course showing test pulse amplitudes from socially isolated mice (black lines depict spaced iTBS). Scattergrams of amplitude and spatial spread outcomes. Donut charts show proportions of different types of response (no change defined as ± 5%). **C.** Example heatmaps for the group-housed response to spaced iTBS. **D.** Time course shows test pulse amplitudes from group housed mice (black lines depict spaced iTBS). Scattergrams of amplitude and spatial spread outcomes, with donut charts showing proportions of different types of response. **E.** Two-way ANOVAs show significant interactions between housing condition and protocol on test pulse amplitude, **F.** and spatial spread. Mean ± SEM; \**P* < 0.05, \*\**P* < 0.01.

By contrast, group-housed mice did not show a significant change in either amplitude (**Figure 4C-D**, *t*(9) = 0.5754, *p* = 0.5791) or spatial spread (*t*(9) = 0.6197, *p* = 0.5508) of test pulse-evoked calcium signals after spaced iTBS induction. This lack of potentiation is consistent with the significant dependence of clinical iTBS plasticity on peak-episode calcium in group-housed mice (**Figure 3E**). For group-housed mice there was a sex difference with the spaced iTBS experiments, with males showing less potentiation to spaced iTBS than females (amplitude, unpaired *t*-test: *t*(8) = 2.795, *p* = 0.0234), but this sex difference was not detected for spatial spread (*t*(8) = 1.640, *p* = 0.1396).

Application of AMPA receptor antagonist NBQX significantly reduced amplitude and spatial spread in both social-isolated (paired *t-*test: amplitude: *t*(9) = 10.48, *p <* 0.0001; spatial spread: *t*(9) = 7.88, *p <* 0.0001) and group-housed mice (paired *t-*test: amplitude: *t*(9) = 10.52, *p <* 0.0001; spatial spread: *t*(9) = 6.92, *p <* 0.0001).

### Synaptic Potentiation by Spaced iTBS Protocol in Socially-Isolated Mice

Two-way ANOVAs investigating the relationships between isolation condition and induction protocol on the outcome of plasticity showed significant interactions for both amplitude (**Figure 4E**, *F*(1, 38) = 7.108, *p* = 0.0112) and spatial spread (**Figure 4F**, *F*(1, 38) = 6.388, *p* = 0.0158). *Post-hoc* Tukey tests reveal that spaced iTBS exerts stronger effects than clinical iTBS on the amplitude (*p* = 0.0394) and spatial spread (*p =* 0.0182) of potentiation in social isolation. This effect is significantly greater than the effect of spaced iTBS on group-housed (amplitude, *p* = 0.1076; spatial spread, *p* = 0.2634). These results indicate that the effectiveness of an iTBS induction protocol to induce potentiation is dependent on whether the mice are group-housed or socially isolated. Specifically, spaced iTBS is highly effective in potentiating test pulse-evoked calcium signals in social isolation.

### Effectiveness of clinical and spaced iTBS in mice that are socially isolated during early adulthood

We have demonstrated that spaced iTBS is more effective at eliciting synaptic potentiation in the prefrontal cortex of socially-isolated mice than clinical iTBS. For this initial experiment, we used *juvenile-onset* social isolation (from P21), which is considered a more extreme behavioural intervention than social isolation in adulthood (Grigoryan et al., 2022; Sun et al., 2017). Next, we asked whether spaced iTBS is effective at eliciting potentiation in *adult-onset* social isolation (from >P60). For these experiments, we isolated adult mice (**Figure 5A**, **Table 1**) and performed either spaced iTBS or clinical iTBS.

**Figure 5:**
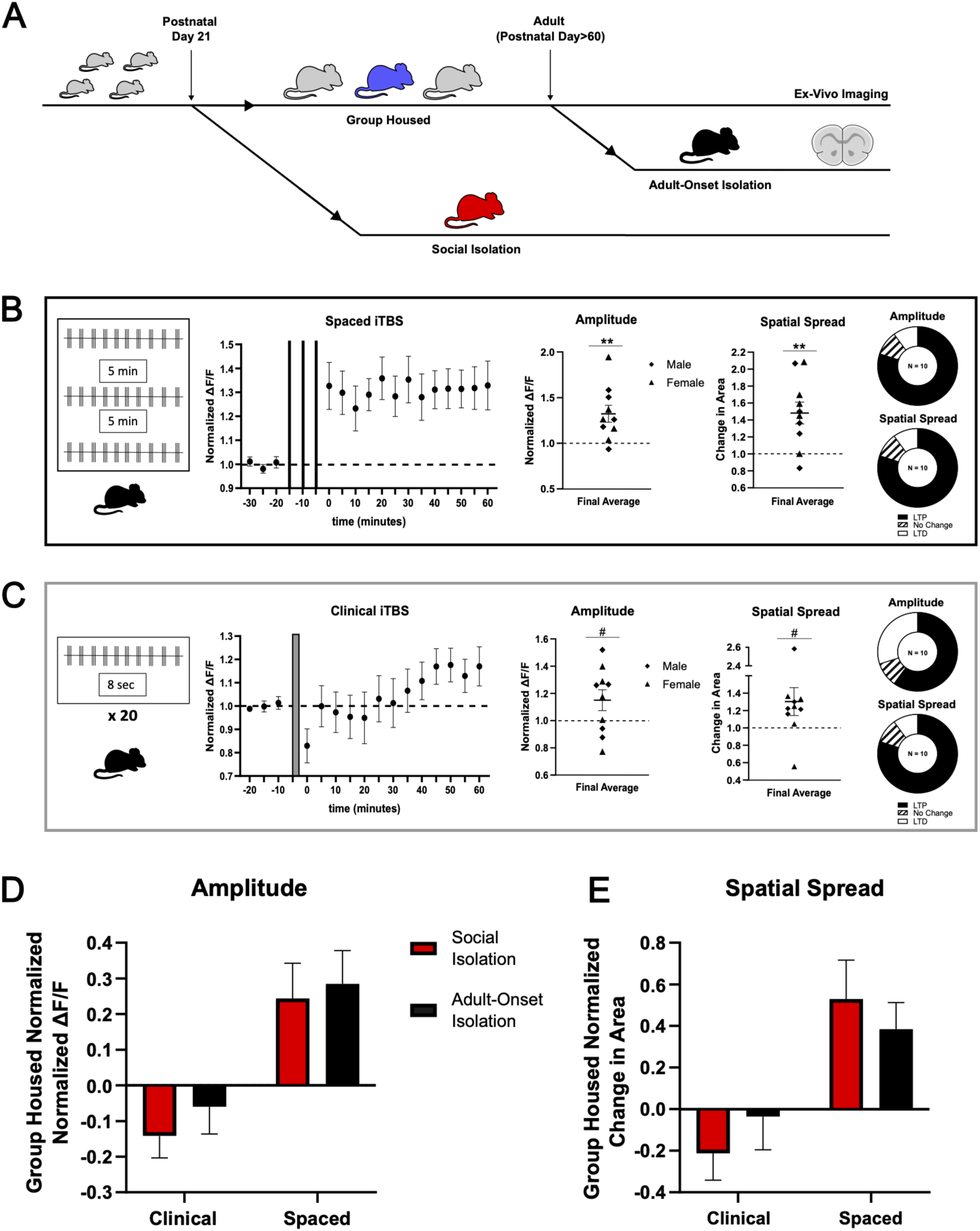
Spaced iTBS is also effective for synaptic potentiation in adult-onset isolation. **A**. Schematic of juvenile-onset social isolation (red mouse) versus adult-onset (black), compared with group-housed controls (blue mouse) as well as timing of *ex vivo* experiments in adulthood. **B.** Schematic of spaced iTBS (left). Time-course shows test pulse-evoked calcium signal amplitudes from adult-onset isolation group (black lines depict the spaced iTBS). Scattergrams of calcium amplitude and spatial spread outcomes. Donut charts show proportions of different types of response (no change defined as ± 5%). **C.** Schematic of clinical iTBS (left). Time-course of calcium amplitudes in adult-onset isolation, (gray bar depicts clinical iTBS). Scattergrams of final calcium amplitude and spatial spread outcomes, with donut charts showing proportions of different types of response. **D.** Illustration of the impact of the different protocols in juvenile- and adult-onset social isolation relative to group-housed controls for amplitude, **E.** and spatial spread. Mean ± SEM; #*P* < 0.1, \*\**P* < 0.01.

In adult-onset isolation, spaced iTBS elicited potentiation, including a significant increase in amplitude (**Figure 5B**; one-sample *t-*test, *t*(9) = 3.479, *p* = 0.0070) and spatial spread (*t*(9) = 3.751, *p* = 0.0045). With clinical iTBS, there were only trends in amplitude (**Figure 5C**; one-sample *t-*test, *t*(9) = 1.962, *p* = 0.0814) and spatial spread (*t*(9) = 1.891, *p* = 0.0911). No significant correlation was detected between age of onset of social isolation and the potentiation outcome for spaced iTBS (Two-way ANOVA, amplitude: *F*(1, 18) = 0.2838, *p* = 0.6007; spatial spread: *F*(1, 18) = 0.0002586, *p* = 0.9872) nor for clinical iTBS (amplitude: *F*(1, 20) = 1.052, *p* = 0.3174; spatial spread: *F*(1, 20) = 0.1389, *p* = 0.7133). Application of AMPA receptor antagonist NBQX significantly reduced the amplitude and spatial spread in both spaced iTBS (paired t-test: amplitude: *t*(9) = 10.15, *p <* 0.0001; spatial spread: *t*(9) = 11.28, *p <* 0.0001) and clinical iTBS (paired *t*-test: amplitude: *t*(9) = 8.74, *p <* 0.0001; spatial spread: *t*(9) 7.84, *p <* 0.0001).

Bringing together the juvenile- and adult-onset social isolation groups yields a signi@icant main effect of protocol for the amplitude of the synaptic outcome (Two-way ANOVA, amplitude: *F*(1, 36) = 4.523, *p* = 0.0404), with a trend for the effect of protocol on spatial spread (*F*(1, 36) = 3.931, *p* = 0.0581; **Suppl Fig S2A,B**). Normalizing the outcomes for juvenile- and adult-onset social isolation to the group-housed controls illustrates and contextualizes the relative impacts of clinical and spaced iTBS for synaptic potentiation (**Figure 5D-E)**.

Taken together, the social isolation experiments reveal that spaced iTBS is effective at eliciting synaptic potentiation in brain slices of prefrontal cortex from adult mice, regardless of the age of onset of the isolation intervention.

## Discussion

We demonstrate that clinical iTBS reliably potentiates the amplitude and spatial spread of test pulse-induced calcium signals in brain slices of the adult mouse prefrontal cortex. This reliability, however, is diminished in socially isolated mice. In the prefrontal cortex of isolated mice, there is a steeper calcium elevation during clinical iTBS induction, which appears to disrupt the typical relationship between peak calcium and synaptic potentiation. A modified induction pattern with fewer episodes of iTBS, spaced further apart, constrains peak calcium and improves the reliability of synaptic potentiation in social isolation. The efficacy of spaced iTBS is seen in both juvenile- and adult-onset social isolation paradigms.

### Social isolation changes outcomes of theta-burst potentiation in prefrontal cortex

Social withdrawal and isolation are prevalent in depression (Liu et al., 2025; Orben et al., 2020; Santini et al., 2020; Xiong et al., 2023) and have been associated with changes in the human prefrontal cortex (Cacioppo & Patrick, 2014; Pizzagalli & Roberts, 2022; Spreng et al., 2020; Vitale & Smith, 2022). In rodents, social isolation is considered a model of chronic stress relevant to depression (Grigoryan et al., 2022; Han et al., 2018; Takatsu-Coleman et al., 2013; Yates et al., 1991; Goh et al., 2024) and associated with cortical disruption, including altered synaptic activity (Li et al., 2022; Liu et al., 2012; Medendorp et al., 2018; Silva-Gómez et al., 2003; Yamamuro et al., 2018). Many of the cortical changes observed in social isolation are relevant for plasticity, such as alterations in inhibitory circuits, NMDA receptors, and G-protein coupled receptor signalling (Bicks et al., 2020; Biro et al., 2023; Gondora et al., 2021; Medendorp et al., 2018; Okamura et al., 2023; Sargin et al., 2020; Ueno et al., 2017; Yamamuro et al., 2018, 2020; Yuan et al., 2026; Zhao et al., 2025). We find that juvenile-onset social isolation decreases the efficacy of clinical iTBS to achieve plasticity in brain slices of the prefrontal cortex. The decreased outcome is broadly consistent with the ∼40% response rate observed in depressed human patients (Blumberger et al., 2018). It is noteworthy that spaced iTBS, with its fewer stimuli overall, proved reliable at strengthening prefrontal connections in both juvenile- and adult-onset social isolation. Both isolation paradigms have similar behavioural consequences, but the former has a more severe impact (Grigoryan et al., 2022; Sun et al., 2017; Han et al., 2018; Ieraci et al., 2016; Takatsu-Coleman et al., 2013).

### Theta-Burst Stimulation Impact on Integrative Activity and Plasticity

There has been increasing interest in resolving network-level activity in response to plasticity paradigms (Khanzada et al., 2025). Calcium signals are considered a major predictive factor in the outcome of plasticity induction (Shouval et al., 2002; Yeung et al., 2004; Nishiyama et al., 2000), consistent with our observations during clinical iTBS in the group-housed mice (**Figure 3**). Increasing data supports a key role for glutamatergic NMDA receptors in the plasticity elicited by clinical iTBS in human patients and healthy control subjects (Cole et al., 2022; DeMayo et al., 2024; Huang et al., 2007). Bursts of stimuli are efficient in priming the ligand- and voltage-dependent NMDA receptors to yield apical dendritic calcium spikes in rodent *ex vivo* experiments, also known as plateau potentials (Antic et al., 2010; Major et al., 2008; Pagès et al., 2021). Such plateau potentials are efficiently propagated to the soma, potentially connecting the firing of the afferents to the firing of their postsynaptic targets to achieve the “fire together, wire together” criteria for inducing synaptic plasticity.

Clinical iTBS provokes a complex pattern of calcium responses in the prefrontal cortex which is recapitulated across experiments. The calcium response from the first episode is constrained compared to that of subsequent episodes, a pattern consistent with field recordings in the hippocampus during theta-burst that has been attributed to suppression of GABA release (Davies et al., 1991; Larson & Munkácsy, 2015). This change allows the excitatory synaptic response to increase over the subsequent episodes of iTBS. However, it must be underscored that the theta-burst paradigms used in this hippocampal research only investigated a small number of episodes.

Here, we show that the initial episodes of clinical iTBS climb to a peak in terms of their calcium elevation. In the group-housed mice, this peak predicts the subsequent plasticity attained. The calcium elevations then plateau before declining during the second half of clinical iTBS. The decrease in per-episode calcium elevation across the later episodes of clinical iTBS may reflect changes in NMDA receptors (Iacobucci & Popescu, 2024), inhibitory circuits (Benali et al., 2011), or neuromodulator release from prefrontal axon terminals, such as serotonin, which is known to suppress theta responses and NMDA receptors (Turi et al., 2026; Wang et al., 2025).

In social isolation, neither the test pulse-evoked calcium responses nor the first episode calcium responses differ from group-housed mice. However, as the induction episodes progress, the calcium dynamics diverge by housing group, resulting in a significant group by episode interaction. Brain slices from the socially-isolated mice show altered calcium elevation compared to group-housed mice. The pattern and timing suggest that social isolation may compromise negative regulatory mechanisms shaping clinical iTBS calcium elevation in brain slices of group housed mice. Alternatively, there may be upregulation of induction calcium in social isolation, such as increased release of calcium from internal stores via potentially stress-sensitized metabotropic glutamate receptors (Nishiyama et al., 2000; Whitaker et al., 2013). There is much scope for future examination of the likely multi-faceted calcium dynamics associated with clinical iTBS.

In spaced iTBS, by contrast, the 5-minute intervals between episodes constrain calcium dynamics during the induction. Spaced paradigms have been explored in the hippocampus where they enhance protein-synthesis-dependent LTP by allowing time for the activation of second messengers, such as PKA, and the insertion of calcium-permeable AMPA receptors (Park et al., 2016, 2021). In the hippocampal literature, comparison protocols keep the number of stimuli constant, but their components differ considerably from clinical iTBS. The research demonstrates that the initial, spaced theta-burst episode induces PKA-dependent insertion of calcium-permeable AMPA receptors into the postsynaptic membrane, creating a temporal window during which subsequent stimulation elicits additional calcium influx through these receptors (Park et al., 2016, 2021). This mechanism promotes local protein synthesis and the synaptic incorporation of calcium-impermeable AMPARs, thereby consolidating persistent LTP. Importantly, this mechanism depends on the interval between stimulation episodes, as the compressed, comparison protocols curtail the protein synthesis-dependent component of LTP (Park et al., 2016, 2021). Since there has been considerable debate over potential differences in the molecular mechanisms of LTP in the prefrontal cortex versus the hippocampus, deeper investigation is needed into the molecular mechanisms of iTBS-based prefrontal cortical plasticity and its perturbation by social isolation and other models relevant to depression.

### Context and Caveats

We have focused on a common pattern of iTBS in clinical use (Blumberger et al., 2018; Bulteau et al., 2022) as well as a spaced iTBS protocol that was modeled on preclinical research (Park et al., 2016, 2021). We used methods that detect the impact of these different protocols on calcium signaling and the prefrontal cortical calcium response to single test pulses. The *Thy1*-GCaMP6f mice have been shown with retrograde labeling to allow calcium imaging in a mixed population of prefrontal cortex output neurons (Muysers et al., 2024). These include intratelencephalic neurons (Muysers et al., 2024), which have recently shown to respond with strong potentiation to accelerated iTBS protocols *in vivo* (Gongwer et al., 2026; Johnson et al., 2026; Salimi & Ramanathan, 2026).

In conducting this work in acute brain slices, we have restricted the recruitment of neuromodulation to the local environment. *In vivo* experiments, by contrast, recruit a more complicated response via long-range feedback circuits from the prefrontal cortex to the dorsal raphe, nucleus basalis, and ventral tegmental area. Here, we employed electrical stimulation in a manner akin to most *ex vivo* investigations of synaptic plasticity. While this approach yields synaptic responses sensitive to AMPA receptor blockade, it is possible that transcranial magnetic stimulation exerts a different constellation of effects. Recent work delivering iTBS *in vivo*, however, demonstrates that similar outcomes can arise despite profound differences in the type of stimulation applied (Gongwer et al., 2026; Johnson et al., 2026; Salimi & Ramanathan, 2026).

By focusing on one session of clinical iTBS, we investigate an intervention known to elicit synaptic potentiation in human and in mouse *in vivo* (Chung et al., 2017; Salimi & Ramanathan, 2026). However, this initial session is typically only the first of many in the treatment of depression. Furthermore, there are many variables in the timing of clinical iTBS, and here we have only changed the interval length, number of episodes, and total stimuli delivered. Considerable scope remains for future research to examine other variables and to probe iTBS mechanisms in the prefrontal cortex.

### Significance and Clinical implications

Since there is increasing attention to accelerated iTBS, with its high number of episodes and spaced-like repetitions, it appears urgent to rigorously investigate how to deliver the basic components most effectively. Our preclinical research suggests that currently used clinical iTBS protocols may be suboptimal for inducing neuroplasticity in the vulnerable prefrontal cortex. We raise the question whether a longer pause between iTBS episodes improves the efficacy of the intervention, allowing greater impact from fewer stimuli. This work builds on a body of preclinical research that may help identify which modifications of the treatment protocol will be most promising. If the goal of theta-burst treatment is to generate LTP, enhancing the recruitment of the prefrontal cortex to regulate mood, then selecting the most efficient iTBS components and optimizing their timing may help accelerate clinical outcome while potentially reducing risks and side effects.

## Methods

### Animals

Animal experiments were approved by the University of Toronto Temerty Faculty of Medicine Animal Care Committee (protocol #20011621) and were conducted in accordance with the guidelines of the Canadian Council on Animal Care. We used Thy1-GCaMP6f transgenic (hemizygous) mice (Jackson Labs, Strain #:025393; RRID:IMSR_JAX:025393) on a C57BL/6J background expressing fluorescent calcium ion indicator in neurons (Dana et al., 2014). We studied the responses of pyramidal neurons of the prefrontal cortex to superficial cortical electrical stimulation. Male and female mice were weaned at postnatal day (P) 21, separated by sex, and group-housed (2-4 mice per cage) in plastic cages with bedding, environmental enrichment, and *ad libitum* access to food and water. A subset of mice was singly housed under a social isolation paradigm, either from weaning age or in adulthood (>P60) by random selection. Experiments included equal numbers of male and female mice, with subjects age-matched in each experiment. Ages of mice in each experiment and further details of social isolation are shown in **Table 1**.

### Widefield calcium imaging and test pulse stimulation

Coronal brain slices of prefrontal cortex (400 µm, Bregma 2.0-0.5).) were prepared in 4°C sucrose artificial cerebrospinal fluid (ACSF) (254 mM sucrose, 10 mM D-glucose, 26 mM NaHCO3, 2 mM CaCl2, 2 mM MgSO4, 3 mM KCl, and 1.25 mM Na2PO4) with a Dosaka linear slicer (SciMedia, Costa Mesa, CA). Slices were transferred to a holding prechamber (AutoMate Scientific, Berkeley, CA) where they recovered for at least 2 h in oxygenated (95% O2, 5% CO2) ACSF (128 mM NaCl, 10 mM D-glucose, 26 mM NaHCO3, 2 mM CaCl2, 2 mM MgSO4, 3 mM KCl, and 1.25 mM Na2PO4) at 30°C before being used for widefield calcium imaging on an Olympus BX61WI microscope (RRID:SCR_023069) in ACSF at 32°C.

Calcium fluorescence images were collected at 5-minute intervals using a Prime BSI express camera (Teledyne) and μManager software (RRID:SCR_000415), and acquired images were binned every 2 pixels. Calcium changes during experiments were visualized with ImageJ (2.9.0/1.53t; Java 1.8.0_322 [64 bit]; RRID:SCR_003070), with imaging of 5 test pulses every 5 minutes.

Electrical stimulation was delivered to superficial cortex through a bipolar stimulating electrode (FHC) connected to a stimulus isolation unit (DS3, Digitimer). The electrical stimulation and the acquisition of images were both controlled by a Multiclamp 700b (Axon, RRID:SCR_018455) via pClamp software (Axon Instruments, RRID:SCR_011323).

Single test pulse imaging compromised 2 s of recording surrounding each 100 µs test pulse, for a total of 5 test pulses. The test stimuli duration was selected to match the pulse duration commonly used in transcranial magnetic stimulation (Spampinato et al., 2023). To replicate the conditions used in traditional plasticity studies, test pulses were delivered at 0.1 Hz for the duration of the experiment.

### Inclusion and exclusion criteria

Each experiment included one brain slice/mouse. Stimulation intensity was adjusted to obtain a calcium signal in response to a single test-pulse that was ∼3x the noise of the baseline recording (mean ± SEM: 80 ± 22 µA) and followed to determine baseline stability. Stimulus intensity versus response curves were assessed across a range of current (20 - 150 µA) for a subgroup of recordings from a matched set of group-housed and socially-isolated mice (**Suppl Fig S1**).

A small number of brain slices were excluded before experiment completion for one of the following reasons: a stable baseline was not achieved (*n* = 3) or a sustained increase in resting calcium signal developed during the experiment, indicating a deterioration in the health of the brain slice (*n* = 3).

### Theta-burst stimulation plasticity induction protocols

Following a 15-minute stable baseline, the slice received either an intermittent theta-burst stimulation for clinical iTBS (burst: 3 pulses at 50 Hz; episode/train: 10 bursts at 5 Hz; paradigm: 20 episodes at 0.1 Hz), a control protocol with no induction stimulation, or spaced iTBS (burst: 3 pulses at 50 Hz; episode/train: 10 bursts at 5 Hz; paradigm: 3 episodes with 5-minute intervals). Note that the 0.1 Hz background test pulse stimulation continued during the spaced iTBS to recapitulate the spaced protocol as described (Park et al., 2018, 2021).

Test pulses were resumed 5 minutes after either the start of the clinical iTBS induction, the start of the final spaced iTBS episode, or control. Calcium fluorescence recordings were taken each 5-minutes thereafter for a total of 60 minutes following induction. After the 60-minute recording, competitive AMPA-receptor antagonist NBQX disodium salt (20 µM; 5-10 minutes; HelloBio product code HB0443).

### Data analysis and photobleaching correction

Collected recording (TIFF format) were imported into MATLAB (Matrix Laboratory, version 2024b; RRID:SCR_001622) for data analysis with automated protocols performed blind to condition. Custom processing code was used to divide pixels into regions of interest (ROIs) forming a 22 by 22 grid (∼100 x 100 µm^2^). After isolation of the recording frames, pixel fluorescence intensity was averaged within ROIs for each frame. A first-degree line was fit to the data within each ROI. This line was then subtracted from the data to correct for photobleaching. The corrected recording was used to calculate a ΔF/F value for each recorded pulse.

### Region of interest selection

To determine which ROIs contained a response to test pulse baseline stimulation, the final baseline recording before induction was used. Each region of interest was averaged with the 8 surrounding regions of interest, such that the trace for a single ROI was representative of the 3x3 grid of ROIs surrounding the ROI being examined. The 3x3 grid trace for each ROI was then assessed for local peaks.

To assess the peak values relative to the regular activity in the slice, the standard deviation of the fluorescence response prior to stimulation was obtained, and a z-score representative of peak size relative to baseline activity for each region of interest was calculated. The top 30% of z-scores was selected, and a two-dimensional convolution was applied to verify that the corresponding ROIs were not outliers. The peak values for these ROIs were then extracted, and the largest value was taken as the ROI of interest. This automatic selection of ROIs was blind to condition, as was the manual quality check was also done to confirm that the selected region was within expected confines, and there were no discrepancies between the manual quality check and the automatic selection of ROIs.

### Time course plots

To observe how the calcium fluorescence changed over time, the peak values were extracted for all time points recorded. Data was normalized to the three baseline recordings.

### Heatmaps and spatial spread of response

To visualize the change in amplitude and spatial spread of the response over time, heatmaps were generated using the final two timepoints under the evaluated condition. They represent a Z-score representative of peak size relative to baseline activity for each ROI and are normalized to the ROI at baseline.

To determine how the response is changing spatially over time, ROIs that contained an amplitude response at least half the maximum (ROI of interest) at baseline were included as part of the spatial response to stimuli. All spatial spread values were normalized to baseline area.

### Calcium signals *during* theta-burst induction

Induction calcium response was calculated from the best ROI as determined previously. Recordings underwent photobleaching correction, and absolute values for area under the curve (AUC) of each episode were measured. All the episode values were normalized to measures of baseline test pulses, unless otherwise stated.

### Statistics

Statistical analyses were performed using GraphPad Prism 10 (version 10.4.1; RRID:SCR_002798). Sample sizes were estimated by power analysis from pilot data with a level of significance of 0.05 and a power of 0.90. While both sexes were included in equal numbers, these analyses may be underpowered and therefore should be considered exploratory. For evaluations of synaptic plasticity outcome, the final two recorded timepoints under the evaluated condition were averaged. To visualize the outcomes in donut plots, no change was defined as ±5%. Greater increases were considered LTP, and greater decreases considered long term depression (LTD).

Mean fluorescence amplitude or mean spatial spread changes between protocols, such as control versus clinical iTBS were calculated using unpaired *t*-tests. Changes from baseline were calculated using one-sample *t*-tests and *F* tests for variance. Mean fluorescence amplitude change in cases where pharmacological agents were added at the end of experiments were compared using paired *t*-tests. Housing condition and stimulation protocol effects and interactions were analyzed by two-way ANOVA with Tukey’s *post hoc* comparisons to assess each iTBS paradigm across isolation conditions and contrast the different paradigms within each condition.

For the analysis of calcium elevation *during* theta-burst induction, clinical iTBS and spaced iTBS calcium signals from the individual episodes were analysed using repeated two-way ANOVA with Geisser-Greenhouse correction, unpaired *t*-tests, and correlation analyses. Slopes were obtained by linear regression of first to peak episode area-under-the-curve values.

## Data availability

The data supporting the findings of this study are available within the article, Table, Figures, and supplementary material (Suppl Fig S1, S2). On final publication, the code will be made publicly available on Github: https://github.com/eklambe/Custom-MATLAB-processing-code-files.

## Acknowledgements

We thank Ms. Yao-Fang Tan for expert technical assistance. We are grateful for the support and encouragement of the INSPiRE-D Study Group (Improving Neuroplasticity through Spaced Prefrontal intermittent-Theta-Burst-Stimulation REfinement in Depression), whose members include: Dr. Aram Abbasian, Ms. Salma Abdelghaffar, Dr. Branka Agic, Dr. Daniel Blumberger, Dr. Heather Brooks, Dr. Mary Chiu, Ms. Dewi Clark, Dr. Graham Collingridge, Dr. Simon Davies, Dr. Peter Finnie, Mr. Norm Fisher, Ms. Maryse Gad, Dr. John Georgiou, Ms. Angel Hsieh, Dr. Rachael Ingram, Dr. Evelyn Lambe, Dr. Clement Ma, Dr. Bernadette Mdawar, Mr. Firdosi Mehta, Dr. Roumen Milev, Dr. Benoit Mulsant, Dr. Vladislav Myrov, Dr. Tarek Rajji, Dr. Thomas Sanderson, Dr. Sanjeev Sockalingam, Ms. Cara Sullivan, Ms. Quincy Vaz, Dr. Sridevi Venkatesan, Dr. Daphne Voineskos, Dr. Albert Wong, Dr. Herbert Yao, Dr. Reza Zomorrodi, Ms. Angela Zolis, Dr. Brigitte Zrenner, Dr. Christoph Zrenner. We thank Dr. Sanjeev Sockalingham of CAMH for help in obtaining funding for this project. We also appreciate the feedback and insightful questions from Dr. Robert Bonin, Dr. Zhong-Ping Feng, Dr. Beverley Orser, and Dr. Etienne Sibille of the Temerty Faculty of Medicine at the University of Toronto.

## Funding

The research was funded by the Bell Let’s Talk-Brain Canada Mental Health Research Program (EKL, GLC, TKR) and by the Canadian Institutes of Health Research (CIHR, PRJ-178372, EKL).

## Conflict of Interest Statement

Dr. Rajji has received research support from Brain Canada, Brain and Behavior Research Foundation, BrightFocus Foundation, Canada Foundation for Innovation, Canada Research Chair, Canadian Institutes of Health Research, Centre for Aging and Brain Health Innovation, National Institutes of Health, Ontario Ministry of Health and Long-Term Care, Ontario Ministry of Research and Innovation, and the Weston Brain Institute. For an investigator-initiated study, Dr. Rajji received in-kind equipment support from Newronika, and in-kind research online accounts from Scientific Brain Training Pro and participated in 2021 and 2022 in an advisory activity for Biogen Canada Inc. In 2023 and 2024, Dr. Rajji was an *ex officio* member of the Board of Trustees of the Centre for Addiction and Mental Health (CAMH) in his role as Chair of the Medical Advisory Committee at CAMH. Dr. Rajji maintains a collaborator scientist and a courtesy appointment at CAMH and a Status-Only appointment at the University of Toronto. Dr. Rajji is also an inventor on the United States Provisional Patent No. 17/396,030 that describes cell-based assays and kits for assessing serum cholinergic receptor activity.

The other authors report no conflicts of interest.

## Contributions

Conceptualization (AZ, AH, SV, RI, JG, CZ, GLC, TKR, EKL); Methodology and Validation (AZ, AH, SV); Investigation (AZ, AH, SV); Formal analysis (AZ, AH); Visualization (AZ, AH); Writing - original draft (AZ, AH, EKL); Writing - review and editing (AZ, AH, SV, RI, JG, CZ, GLC, TKR, EKL); Supervision (SV, EKL); Resources (GLC, TKR, EKL)

## Notes

### Competing Interest Statement

The authors report no conflicts of interest, except for the following. Dr. Rajji has received research support from Brain Canada, Brain and Behavior Research Foundation, BrightFocus Foundation, Canada Foundation for Innovation, Canada Research Chair, Canadian Institutes of Health Research, Centre for Aging and Brain Health Innovation, National Institutes of Health, Ontario Ministry of Health and Long-Term Care, Ontario Ministry of Research and Innovation, and the Weston Brain Institute. For an investigator-initiated study, Dr. Rajji received in-kind equipment support from Newronika, and in-kind research online accounts from Scientific Brain Training Pro and participated in 2021 and 2022 in an advisory activity for Biogen Canada Inc. In 2023 and 2024, Dr. Rajji was an ex officio member of the Board of Trustees of the Centre for Addiction and Mental Health (CAMH) in his role as Chair of the Medical Advisory Committee at CAMH. Dr. Rajji maintains a collaborator scientist and a courtesy appointment at CAMH and a Status-Only appointment at the University of Toronto. Dr. Rajji is also an inventor on the United States Provisional Patent No. 17/396,030 that describes cell-based assays and kits for assessing serum cholinergic receptor activity.

### Summary of Updates

The manuscript has been updated to clarify the methods and the results. Figures 3 and 5 have been updated to clarify the statistical analyses and the units on the axes. For clarity, the Table has been moved from the Supplementary section to the body of the manuscript.

