## Supplementary material for "Spacing theta-burst stimulation enhances synaptic potentiation in the vulnerable prefrontal cortex": Suppl Fig S1

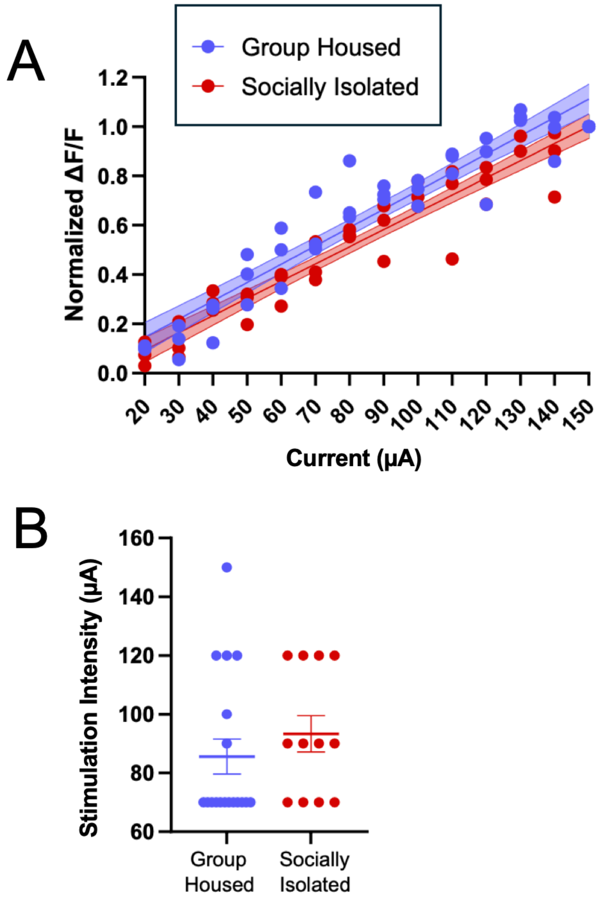


**Supplementary Figure S1: Test pulse stimulus responses by housing condition.**

**A.** Input-output responses for a subgroup of mice (*n* = 6): group housed (blue) and socially isolated (red) Data are normalized to the response at 150 μA. **B.** Stimulation intensity to meet test pulse criteria (~3x baseline noise) did not differ between group housed and socially isolated mice (*n* = 22 mice).
