## Supplementary material for "Spacing theta-burst stimulation enhances synaptic potentiation in the vulnerable prefrontal cortex": Suppl Fig S2

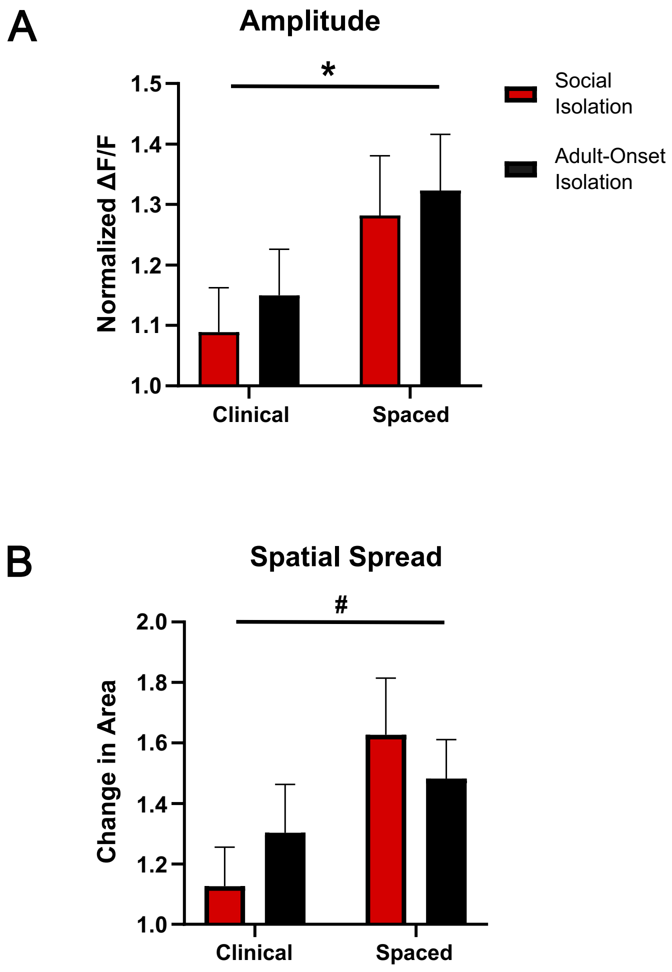


**Supplementary Figure S2: Impact of theta-burst protocol across juvenile- and adult-onset social isolation groups.** **A**. Graph compares the amplitude synaptic outcomes of clinical (600 stimuli, 3 minutes) and spaced (90 stimuli, 10 minutes) induction protocols in juvenile- (red) and adult-onset (black) social isolation. Two-way ANOVA detected a main effect of protocol on the amplitude outcome. **B.** Graph compares the spatial spread synaptic outcomes of these protocols on the same social isolation groups. Two-way ANOVA detected only a trend toward a main effect of protocol on spatial spread outcome. Mean ± SEM; #$P < 0.1$, *$P < 0.05$.
